# Fundamentals of biofilm formation in soil: From functionalized self-assembled monolayers to rewilding

**DOI:** 10.1101/2023.01.30.526320

**Authors:** Saghar Hendiani, Mads Frederik Hansen, Ioannis Kontopoulos, Taru Verma, Milda Pucetaite, Mette Burmølle, Madeleine Ramstedt, Karina Krarup Svenninggaard Sand

## Abstract

Surface energy and surface charges play crucial roles in bacterial adhesion and biofilm formation, however the mechanisms underlying the bacteria-surface interaction, particularly on the formation of soil biofilms, remain unclear. In spite of considering the spatiotemporal dynamics of biofilm formation on different soil surfaces, we compared the impact of four different substrates on bacterial attachment and biofilm formation. The substrates were constituted of gold layer covered by NH_2_^+^, CH_3_, COO^-^ and OH-terminated self-assembled monolayers (SAMS). Two soil habitat bacteria with different Gram barriers, *Bacillus subtilis* and *Acinetobacter baylyi*, were grown with incubation times of 6-72 h on each type of surfaces. Bacterial attachment and biofilm formation was assessed using metabolic activity of the cells adhered to the surfaces. The spatial distribution of adhere bacteria was visualized by scanning electron microscopy and confocal laser scanning microscopy. We also investigated whether the surface impacts the biofilm matrix composition. A general view of our results suggests a major influence of the surface chemistry on bacterial potential to form biofilms. The hydrophobic or positively charged substrates attract bacteria while a lack of attachment and biofilm formation on hydrophilic and negatively charged surfaces. This work points out the potential of surface treatments in the environment where it is intended to either repel or attract bacteria.

## Introduction

Many bacteria live as multicellular communities encapsulated within a self-produced matrix of extracellular polymeric substances. These communities, named biofilms, can be free floating aggregates, but do often adhere to and develop on surfaces [1]. Over time, biofilm cells divide and secrete matrix components such as polysaccharides, proteins, lipids, and nucleic acids. The composition of biopolymers is dependent on the species present, and specific organisms secrete different components dependent on cues and conditions [2]. The biofilm community enable emergent properties unique for this lifestyle compared to planktonic growth; once a biofilm reaches a certain state it can exclude foreign invaders [3], generate local niches that maintain species diversity within [4], and create spatial refugees, where cells are protected from viral infections [5]. The matrix also enhances retention of water, enzymes, protects against desiccation and generates a stable environment in harsh, changing surroundings [6, 7].

From a clinical perspective, biofilm formation can prove an immense challenge because of the protective nature of the matrix and since many of the cells enter a metabolic state, where many antibiotics are ineffective [8]. The formation of biofilm can however also be beneficial. In agriculture, a healthy composition of species can prevent invasion of phytopathogens, increase drought tolerance and influence nutrient sorption and processing [9, 10]. Thus, although soil microbial diversity is sensitive to changes in environmental transitions, it is also a facilitator for ecosystem change. It has been proposed that microbial communities are not only indicators of ecosystem health but able to enhance recovery of degraded ecosystems by manipulation [11]. A better understanding of the parameters that control the various stages of biofilm formation thus might open avenues for rewilding and restoration strategies.

The initial step of surface-associated biofilms is adhesion of a few or even just a single bacterium to a surface. How well the organism colonizes is dependent on the characteristics, appendages and chemical properties of the cell [12], but also the environmental conditions e,g., temperature, pH or mechanical forces [13, 14], and surface properties of the substrate [15]. The very first encounter between cell and surface is primarily influenced by interfacial electrostatic and van der Waals forces. The interaction between the bacterial cell wall and the surface can be attractive, repulsive or nonspecific [16]. Lorite et al. [17] discovered that culture media was able to create a thin film that conditioned the surface and improved the attachment by decreasing hydrophilicity. Others have since then also observed increased biofilm formation when hydrophobic surfaces were applied [18, 19]. There are however other studies, such as Hong et al. [20], which found that bacterial affinity to minerals was conditioned by electrical property rather than hydrophobicity.

Subsequent to fast initial reversible interactions, the irreversible attachment process often occurs on a time scale of several hours. The irreversible attachment is thought to involve hydrophobic interactions between the hydrophobic region of the cell appendages and the surface and in this step, adhesins such as curli, fimbriae, flagella and pili manifest the transition from reversible to irreversible cell attachment [15]. In general, the binding potential is a function of surface properties, the bacterial cell and how well they interact under given conditions. The sum of interactions determines whether a surface attracts or repels a bacterium. The specific factors that govern microbial attachment to surfaces are however not yet fully understood nor is the temporal interplay with the surface.

The properties of biofilms change as the community matures. Cells specialize and adjust expression patterns according to position and local conditions [21, 22]. Eventually, some cells might disperse as a response to certain cues e.g., starvation or quorum-sensing signaling [23, 24]. This current understanding of the spatiotemporal dynamics of biofilm formation is based on studies on common lab surfaces such as agar, glass, or plastic. In the present study and in regard to offering the potential of biofilms for rewilding purposes, we investigated how substrate chemistry affected the various stages of biofilm development. We characterized structure, matrix composition and metabolism at various time points on various surfaces with different surface energy and charge. Specifically, we used, quantification of triphenyl tetrazolium chloride (TTC) conversion, Scanning Electron Microscopy (SEM), Confocal Laser Scanning Microscopy (CLSM), and Micro-Fourier Transform Infrared Spectroscopy (Micro-FTIR) to address these questions. To generate different substrates with different properties we used self-assembled monolayers (SAMS) of alkanethiol terminated with CH_3_, OH, COO^-^, and NH_2_^+^, respectively. This represents surfaces with hydrophobic, hydrophilic, negative, and positive charge features, respectively. Biofilms were formed by *Acinetobacter baylyi* and *Bacillus subtilis*, to evaluate differences between Gram-negative and Gram-positive cells.

## Results

### Bacterial attachment and biofilm formation are substrate-dependent

In order to monitor the temporal biofilm dynamic, we measured the conversion of TTC at six different time points. TTC conversion is a proxy of metabolic activity, as active cells convert this compound to a colored formazan derivative that can be quantified as color change from yellow to red. Since it is reasonable to assume that the vast majority of cells are active at the early stages of biofilm formation [25], the TTC conversion also indicates the initial biomass present on the various surfaces tested. With this assay, we observed a peak in activity after 24 hours independent of bacterial species and surface substrate (Fig. 1). For both species, the CH_3_ and NH_2_^+^ treatment (hydrophobic and positively charged, respectively) yielded higher activity than the OH and COO^-^ treatments (T=24h, P<0.05). The difference between the two attractive surfaces (CH_3_ and NH_2_^+^) was significant for *B. subtilis* (T=24h, P=0.02) but not for *A. baylyi* (T=24h, P=0.35). The difference between the two less attractive substrates (OH and COO^-^) was not significant for either of the two species (T=24h, P>0.05).

**Fig 1.**
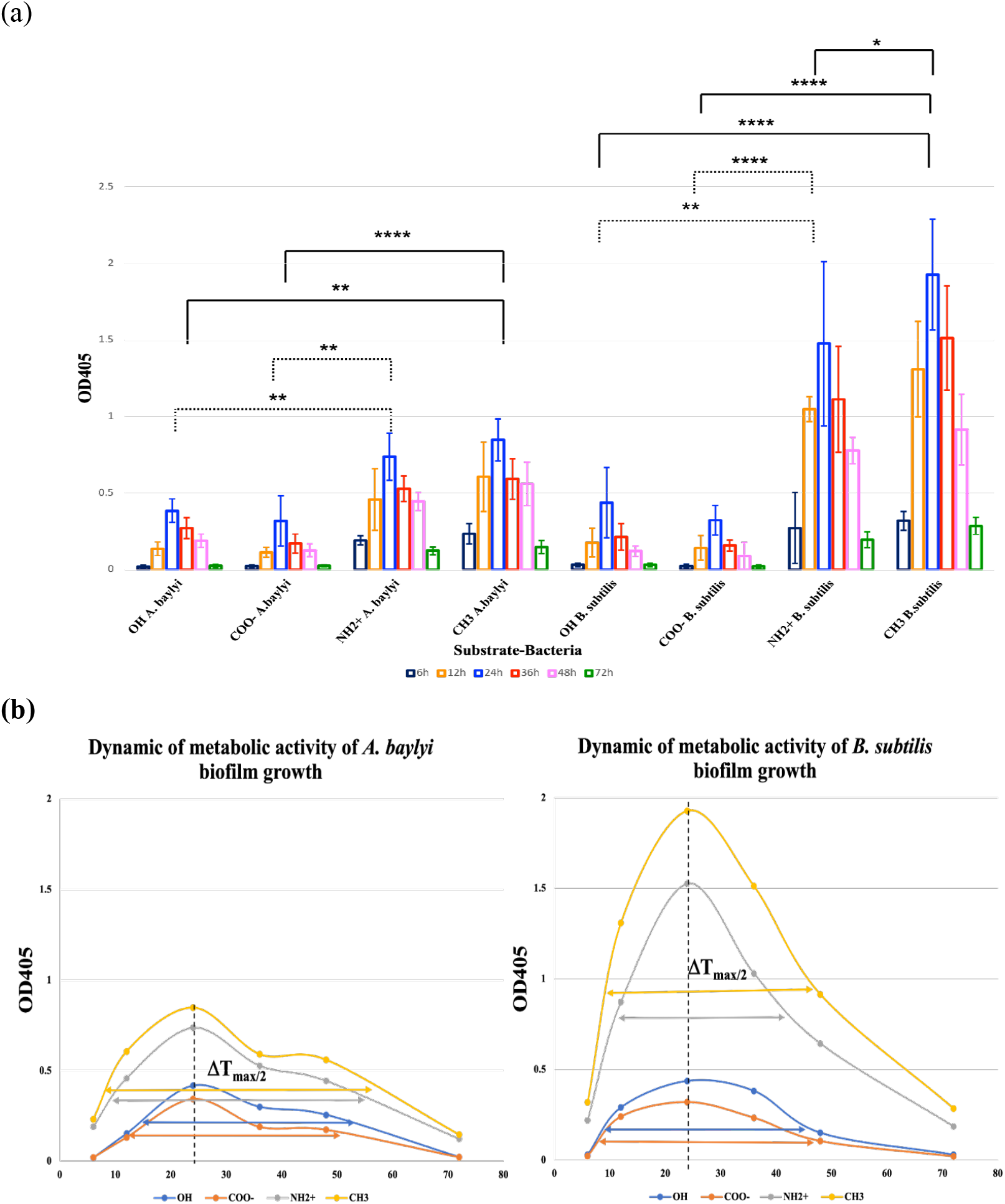
Bacterial metabolism is surface dependent. The TTC conversion was measured for *A. baylyi* and *B. subtilis* biofilms grown on substrates of OH, COO^-^, NH_2_^+^ and CH_3_, respectively, after 6, 12, 24, 36, 48 and 72 hours. TTC-converted formazan derivatives were measured at low frequency wavelength (405nm) and indicate metabolic activity. a) metabolic activity of bacteria at specific time points, b) Dynamics of metabolic activity of biofilms grown on OH, COO^-^, NH_2_^+^ and CH_3_-substrates. Cycle duration is defined by FWHM at △T_max/2_. *Acinetobacter baylyi* exhibit the highest level of activity at 24h post inoculation independent of surface substrates. The hydrophobic and positively charged surfaces (CH_3_ and NH_2_^+^) did however yield a significantly higher activity than the hydrophilic and negatively charged substrates (OH and COO^-^). After 24 h, the activity drops, which can be due to reduced activity or reduced number of cells as results of biofilm dispersal. The same trend was observed for *Bacillus subtilis*. This strain did however reach a higher level of activity and exhibited a significant difference between CH_3_ and NH_2_^+^ after 24 hours with the maximum activity reached on a hydrophobic CH_3_ substrate. The data are representative of six biological replicates and are plotted as mean ±SD. * represents p<0.05; ** p<0.01; *** p<0.001 and **** p<0.0001.

The temporal dynamics of activity of both bacteria exhibit a bell-shaped diagram independent of substrate (Fig. 1.b). *A. baylyi* do however seem to reach a plateau at 36h with a stable activity until 48h where a sudden drop occurs. This trend is independent of substrate. The same pattern is not observed for *B. subtilis*, where the negative rate in activity is reduced after 48h. The final measurement is however as low as the initial measurement at 6h, which reasonably can be regarded as the minimum level detectable. Thus, this minimum could potentially have been reached at an earlier time point than 72h, and hence the final slope of activity might be an artifact of the relatively large time interval from 48h to 72h. Results also show that the positions of cycles maxima and minima, as well as widths of cycles waves are substrate-dependent, leading to a faster biomass evolution on CH_3_ and NH_2_^+^-substrates. Table 1. reports the graphical definitions of incubation time of maximal adhesion and the full width-at-half-maximum (FWHM). The strong difference between the FWHM values, but the same incubation times at the maxima of adhesion, point out the strong difference in biomass evolution for all four substrates.

**Table 1.** Graphically determined positions and widths of adhesion waves on substrates Structure of biofilms are specific for each strain on each surface.

| Bacteria | Substrate | Incubation time at maximal adhesion,<br>*T*<sub>max</sub> (h) | FWHM at *T*<sub>max</sub>/2<br>(h) |
| --- | --- | --- | --- |
| *A. baylyi* | OH | 24 | 33 |
|  | COO<sup>-</sup> | 24 | 23 |
|  | NH<sub>2</sub><sup>+</sup> | 24 | 44 |
|  | CH<sub>3</sub> | 24 | 45 |
| *B. subtilis* | OH | 24 | 22 |
|  | COO<sup>-</sup> | 24 | 21 |
|  | NH<sub>2</sub><sup>+</sup> | 24 | 42 |
|  | CH<sub>3</sub> | 24 | 46 |

In order to clarify whether the metabolic activity reflected the present biomass on the substrates, we visualized the biofilms on the respective surfaces at different time points. Initially we applied scanning electron microscopy (SEM) to visualize biofilm formation after 24 hours of incubation. These images verified a higher biomass on NH_2_^+^ and CH_3_ substrates for both bacteria. Thus, the maximum metabolic activity at 24 hours (Fig. 1) is likely due to a higher number of cells attached. The high resolution of SEM also enabled identification of the EPS matrix uniquely for these two substrates. No matrix was observed for the OH and COO^-^ substrates and the distribution of bacteria on OH and COO^-^ substrates were not uniform and scattered attachment of individual bacterial cells was obvious (Fig. 2). According to Fig. 2 three-dimensional biofilm structure is visible for *B. subtilis* on NH_2_^+^ and CH_3_-substrate and for *A. baylyi* on CH_3_ whereas the tendency to form bacterial monolayer was observed for *A. baylyi* on NH_2_^+^. Interestingly, morphological differences are observed between the structure of biofilms formed on NH_2_^+^ and CH_3_. Uniform biofilm structure and the highest amount of EPS was produced in contact with CH_3_-SAMS for both strains. Whereas, the extracellular material detected on *B. subtilis* biofilms on NH_2_^+^ appeared to be composed of tangled thread-like strands forming a complex network. Interestingly, we also observed smaller pearl-looking structures in the biofilms of *B. subtilis* on NH_2_^+^ and CH_3_, indicating that these surfaces enable the formation and entanglement of spores (Fig. 2). On the other hand, bacterial cell elongation (bacilli shape instead of coccobacilli) was observed within the *A. baylyi* monolayer on NH_2_^+^.

**Fig 2.**
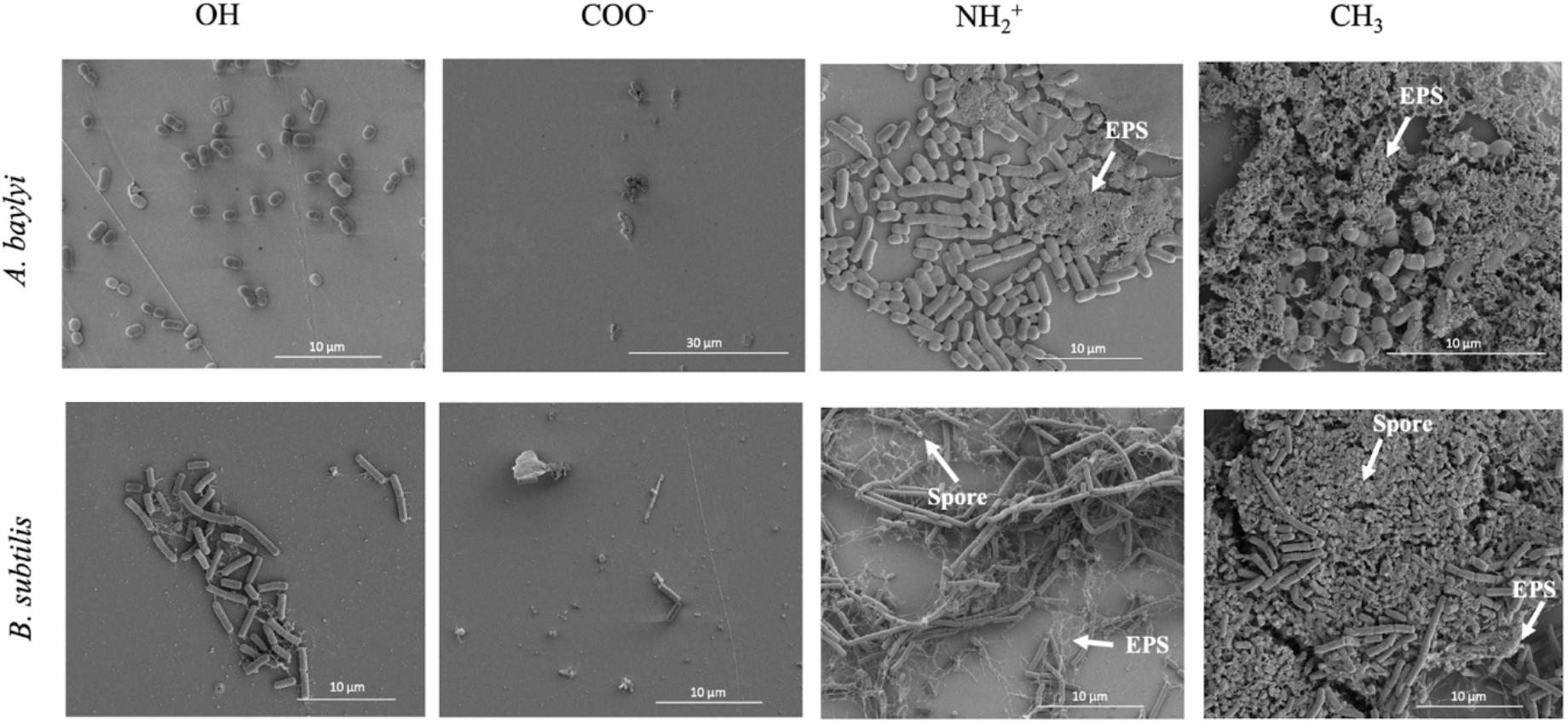
Biomass and matrix production is surface dependent. With the application of scanning electron microscopy the biofilm size and structure was evaluated after 24hours of incubation on various substrates (OH, COO^-^, NH_2_^+^ and CH_3_). On the neutral and negatively charged hydrophilic surfaces (OH and COO^-^, respectively) only few bacteria were present and no matrix could be found. In contrast, the positively charged and hydrophobic surfaces (NH_2_^+^ and CH_3_, respectively) enabled multiple cells to attach and form a biofilm matrix (indicated by EPS arrows). In addition, the two attractive surfaces also enabled the formation of spores integrated within the *Bacillus subtilis* biofilm.

It is necessary to mention that as the dehydration process changes the structure of EPS, we are not able to see the original structure of EPS that is formed by the bacteria, thus, fibrous strand-like structures could be collapsed remnants formed by drying hydrated 3-dimensional structures.

The drop in activity after 24 hours (Fig. 1) can be either due to dispersal of cells from the biofilm or lower metabolism induced by starvation or stress (22-24). With the use of confocal laser scanning microscopy (CLSM) and DNA stains that enable visualization of cells, we monitored temporal biomass development. As shown in Fig. 3, after 3h no remarkable attachment was observed for COO^-^ and OH but the highest number of cells attached to the positively charged NH_2_^+^ substrate. 24h biofilms of A. baylyi on NH_2_^+^ showed monolayer structures whereas their biofilms formed on CH_3_ were more extensive and formed branched arms of multilayer cells. On the contrary, B. subtilis cells assembled in clusters on NH_2_^+^ and CH_3_. The images indicated that the drop of activity after 24 hours was a result of dispersal. We were able to image most biomass at 24 hours independent of substrate (Fig. 3).

**Fig 3.**
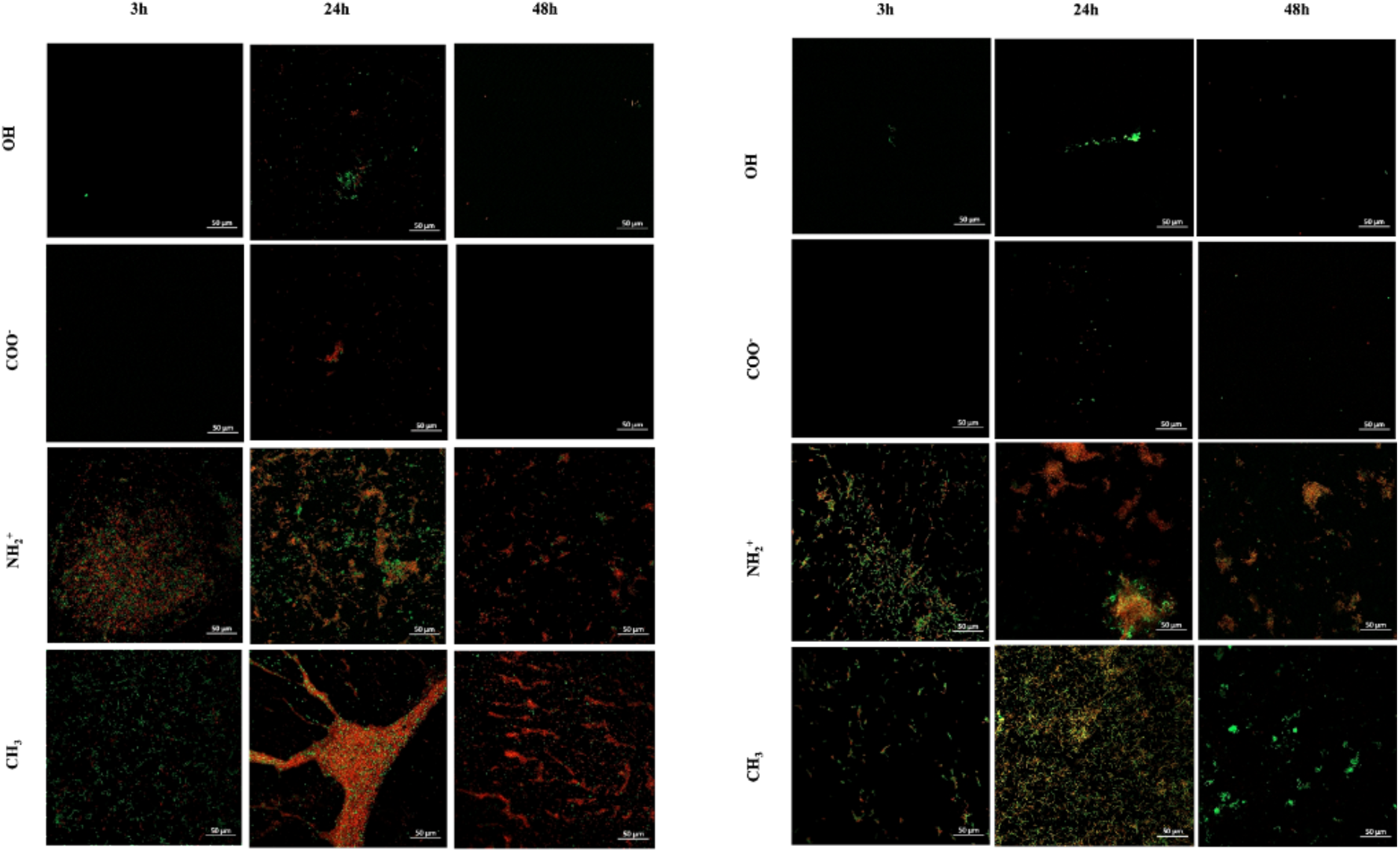
Cells disperse on all substrates tested. The use of two distinctive DNA stains enable visualization of biomass. The cell permeable SYTO9 stains all DNA present (green) including that in living cells, while the cell impermeable propidium iodide (red) stains compromised cells and free DNA present. After staining cells, images of biomass were acquired by confocal laser scanning microscopy. These images showed a higher amount of biomass at 24 hours, whereas the time points earlier and later (3h and 48h, respectively) less cells were visible for all substrates tested. This indicates that cells of both *Acinetobacter baylyi* (left) and *Bacillus subtilis* (right) disperse at some point after 24h. The NH_2_^+^ and CH_3_ treated surfaces enabled higher biomass development than the OH and COO^-^ treated surfaces. These results are in line with observations of maximum activity on these substrates and verify that the elevated activity is biomass related. Images a representative of multiple images acquired and solid lines indicate 50um.

A puzzling observation on the CLSM images was a relatively large amount of propidium iodide staining at 24h and later. This stain is often referred to as the dead stain, as it cannot penetrate intact cells, but rather stain compromised cells and free nucleic acids. Especially *A. baylyi* on CH_3_ substrate at 24 and 48 hours bound a relatively large amount of this stain (Fig. 3). This could indicate that either 1) a large amount of cells is dead at this time point or 2) nucleic acids are a pivotal component of the *Acinetobacter* biofilm. Based on the activity measurement, where activity peaked at 24 h (Fig. 1) it would seem counterintuitive that cells would be dead at this hour.

### No changes in chemical components of biofilm cells

After establishing that hydrophobic and positively charged substrates were more attractive than hydrophilic and negatively charged substrates, we investigated whether the substrate influenced matrix composition. In order to do so, Micro-Fourier Transform Infrared Spectroscopy (Micro-FTIR) analyzed biofilms from three different time points on all substrates. The infrared spectra recorded in transflection mode correspond to the presence of matrix components i.e. proteins, polysaccharides, phospholipids and nucleic acids.The assignment of IR absorbance spectral bands to the functional groups are presented in Fig. 4. Interestingly, neither the shape nor the peak position varied from the various conditions. Only the absorbance intensity varied between samples, with the highest peaks present at 24h (Fig. 4). Spectral signatures indicating presence of cells/EPS appear at the onset of biofilm formation (at 6h) for both strains grown on NH_2_^+^ and CH_3_, while they appear at 24 h for the ones grown on OH and COO^-^substrates. The amount of proteins, nucleic acids, lipopeptide, polysaccharides and phospholipids reached the peak at 24 h for all the substrates, followed by a decrease with biofilm dispersal at 48h. The changes of exopolysaccharides and proteins in biofilm were consistent with the variation of biofilm formation stages.

**Fig 4.**
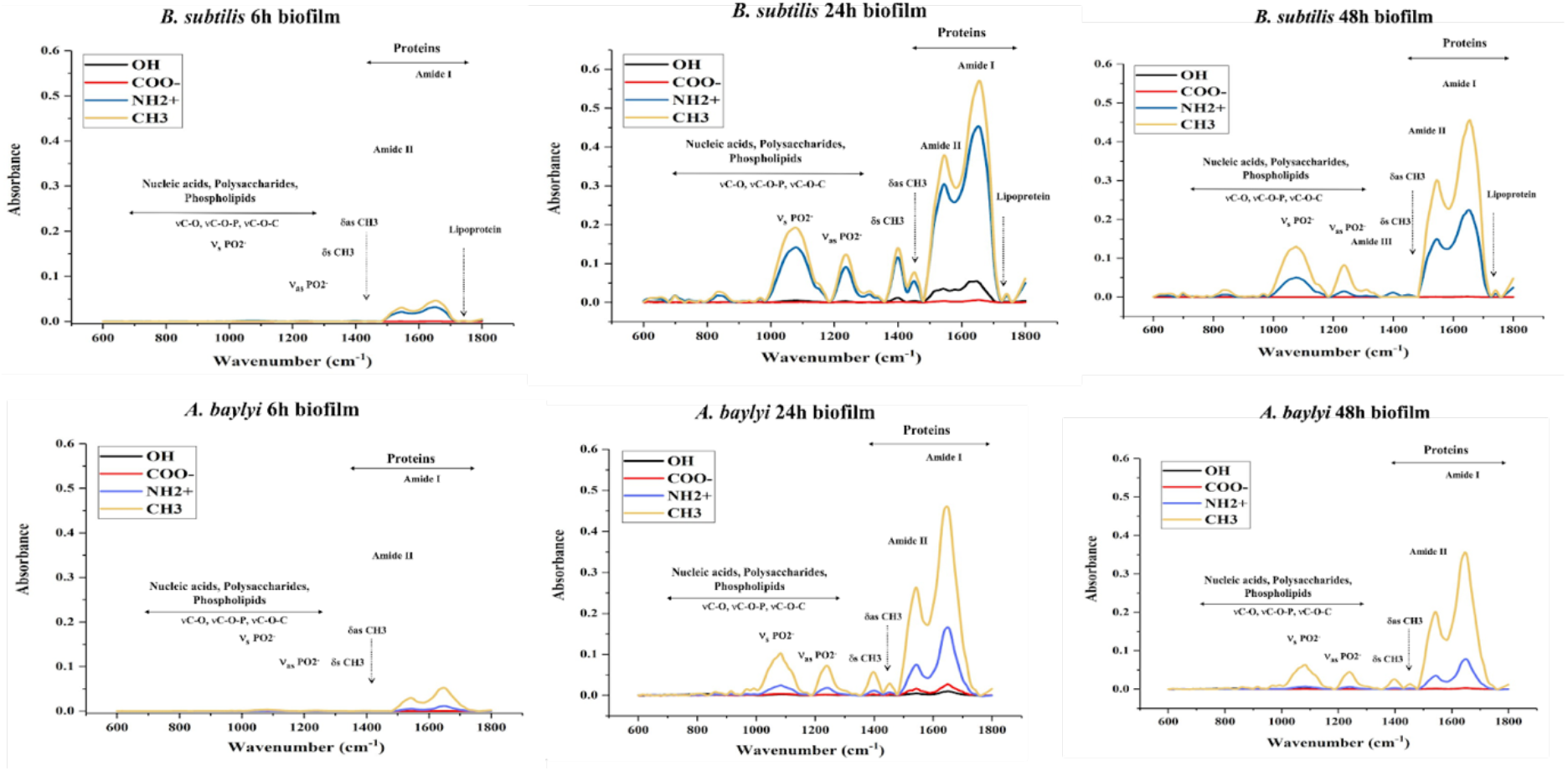
Different substrates enable different levels of matrix components but do not induce differential matrix compositions. The temporal chemical composition of the biofilm matrix was evaluated by Micro-FTIR. Few and small peaks are present for CH_3_ (yellow) and NH_2_^+^ (blue) substrates at 6h biofilms at the high end of the spectra. These peaks increase over time and peaks also emerge at lower wavelengths. The peaks are highest independent of bacterial species at 24h. The reduction in intensity at 48h indicates that less biofilm matrix and cells are present, and hence that biofilm-associated cells have dispersed.

Infrared spectra can reveal changes in relative orientation and hydrogen bonding of the peptide C=O groups, which are the main structure-related oscillators. A detailed second derivative analysis of the 1750-1500 cm^-1^ area offers significant advantages in order to identify and characterize the main component bands underneath the amide I band in terms of position and intensity. Such identification allows the assessment of the secondary structure of proteins.

The spectra of both the *A. baylyi* and *B.subtilis* are dominated by the amide I and amide II protein bands positioned at approx. 1650 cm^-1^ and 1545 cm^-1^ and mainly originating from the C=O stretching and N-H bending vibrations respectively. The second derivative spectra revealed that the amide I band is further composed of two components: one at around 1650 cm^-1^ linked to α-helix structure of protein, and second at around 1630 cm^-1^ associated with the β-sheet structure of protein. For the *A. baylyi* biofilms, small shifts to these bands are observed in the spectra of the biofilms grown on the different substrate and at different time points of biofilm formation. Specifically, for CH_3_ and NH2 substrates at 6 h, the α-helix band is positioned at 1653 cm^-1^, then shifts to 1654 cm^-1^ at 24 h and returns to 1653 cm^- 1^ at 48 h. The β-sheet band shifts in the same time pattern, from 1632 cm^-1^ to 1634 cm^-1^ and back to 1632 cm^-1^ on the CH_3_ substrate, and from 1635 cm^-1^ to 1637 cm^-1^ and back to 1635 cm^-1^ on the NH2 substrate. On the OH substrate, the respective shifts take place from 1653 cm^-1^ to 1654 cm^-1^ and back to 1653 cm^-1^, and from 1634 cm^-1^ to 1636 cm^-1^ and back to 1634 cm^-1^. Lastly, for the COOH substrate, the bands do not shift in time, but are positioned at 1651 cm^-1^ and 1625 cm^-1^ respectively. No band shifts are observed for the amide II band, but the peak position differs depending on the substrate the biofilm was grown on: for the CH_3_, NH2 and COOH substrates it is at 1545 cm^-1^, while for the OH – at 1547 cm^-1^.

For *B. subtilis* biofilms, the amide I band shifts are observed for the CH_3_ and NH2 substrates: from 1652 cm^-1^ at 6 h to 1657 cm^-1^ at 24 h and back to 1652 cm^-1^ at 48 h for the α-helix band; and from 1631 cm^- 1^ to 1634 cm^-1^ and back to 1631 cm^-1^ for the β-sheet band. No shifts are observed for the OH and COOH substrates. However, for the OH substrate, the α-helix band is positioned at 1651 cm^-1^, the β-sheet band, which also displays much higher intensity – at 1623 cm^-1^, and the amide II – at 1541 cm^-1^. Spectral bands at 1577 cm^-1^ and 1513 cm^-1^ assigned to ring vibrations in aromatic compounds are more intense in these samples as well. Meanwhile, for the COOH substrate, the α-helix band is positioned at 1655 cm^-1^, the β-sheet band is split and has two components at 1638 cm^-1^ and 1628 cm^-1^, and the amide II – at 1546 cm^-1^.

Amide I/II values also support differences in secondary structure as seen in the second derivative spectra.

**Table 2.** Amide I / Amide II ratio for each bacteria on each surface-time.

| **Table 2. Amide I / Amide II ratio for each bacteria on each surface-time** |  |  |
| --- | --- | --- |
|  | *A. baylyi* | *B. subtilis* |
| **CH<sub>3</sub>-6h** | 1.77 | 1.51 |
| **CH<sub>3</sub>-24h** | 1.75 | 1.51 |
| **CH<sub>3</sub>-48h** | 1.77 | 1.51 |
| **NH<sub>2</sub>-6h** | 2.22 | 1.50 |
| **NH<sub>2</sub>-24h** | 2.20 | 1.49 |
| **NH<sub>2</sub>-48h** | 2.23 | 1.49 |
| **COOH-6h** | 1.69 | 1.91 |
| **COOH-24h** | 1.67 | 1.90 |
| **COOH-48h** | 1.70 | 1.91 |
| **OH-6h** | 2.08 | 1.58 |
| **OH-24h** | 2.06 | 1.56 |
| **OH-48h** | 2.08 | 1.58 |

## Discussion

Rewilding is the process of recycling and improving unhealthy soil by deletion of previously used fertilizers, herbicides and pesticides and returning the useful microorganism communities back. Microbial biofilms play a major role in rewilding by forming food webs and ensuring nutrient cycles [26], bioremediate the natural systems [27], driving the plant protection, health and productivity under a changing climate [28]. On the other hand, biofilms can promote survival sites for the persistence of plants, animals and human pathogens, produce harmful toxins, erode soil surfaces in natural as well as man-made settings; all of which have notable health and economic consequences [26]. An initial prevention of bacterial attachment would reduce cost and labor associated with eradication of biofilms. This could be example be in instruments used at slaughterhouses or in medical implants, infections often are associated with insertion of a contaminated foreign body [29, 30]. The potential of using modified surfaces to push bacterial cells towards forming biofilms or inhibit their initial attachment can be used in many applications. Treated surfaces can also be used where biofilm formation is wanted.

In order to understand the fundamentals of biofilm formation and extrapolate it to soil environment for rewilding, we have utilized four gold-coated functionalized silicon wafer surfaces consisting of: hydrophobic surfaces with long-chain hydrocarbons (1-undecanethiol), hydrophilic surfaces with negative zeta potential value (1-Mercaptoundecanoic acid), hydrophilic surfaces with positive zeta potential value (1-amino-1-undecanthiol hydrochloride), and hydrophilic surfaces with neutral zeta potential value (1-Mercapto-1-undecanol). The surfaces had OH, COO^-^, NH_2_^+^ and CH_3_ terminated groups, respectively. Self-assembled monolayers on silicon wafers are considered a good method to create well-defined smooth surfaces [31] allowing for direct comparison of the influence of different headgroup chemistry. These indicated that the substrates were physically the same and chemically well-defined and thus ready to act like a probe to study bacterial adsorption and biofilm formation fundamental mechanisms.

In the beginning of biofilm formation, The formation of the conditioning film happens when macromolecules from environmental media adsorb to the substrate [17]. It is assumed that protein adsorption and bacterial adhesion to surfaces follow the same rules, thus, a surface preventing nonspecific protein adsorption is assumed to some extent also resist bacterial adhesion and subsequent biofilm formation (Cheng, G. et al., 2007). Cunliffe, D., et al 1999 also showed that hydrophilic uncharged surfaces prevent protein adsorption and cell attachment at highest levels [32]. On the other hand, Ostuni, et al. concluded that there is very little or no correlation between protein adsorption and cell adsorption to a surface [33]. In order to investigate if surface preconditioning from growth media would play a role in this system, surfaces incubated with growth media for 24 hours were analyzed for surface chemical composition using XPS. The results show that the functionalized surfaces adsorbed very low amounts of substances from the nutritive media. The amount of adsorbed substances appeared to differ between culture media. This most likely reflected their differences in chemical composition as well as more physicochemical characteristics such as ionic strength etc. However, overall the adsorption was too low to enable ranking our surfaces as more or less efficient in preventing adhesion of substances from growth medium (see supplemental information).

Regarding the quantity of bacterial attachment and biofilm formation based on metabolic activity assay, our study shows a significant difference between bacterial tendency to attach to hydrophobic surface with CH_3_ and hydrophilic positive surface with NH_2_^+^ compared to hydrophilic uncharged substrate with OH and hydrophilic negative substrate with COO^-^ for all the incubation times. Evidence shows that bacteria with hydrophobic cell walls prefer hydrophobic substrates [34]. Several determinants are involved in bacterial-surface interactions such as surface characteristics, bacterial characteristics (cell wall and cell appendages like pili, lipopolysaccharide (LPS), outer membrane proteins (OMP)), nutrients, cell density, and time of exposure [35]. Both strains studied here present hydrophobic properties on their cell wall and cell appendages which can explain their attraction towards surfaces with CH_3_. Gohl, et al. demonstrated that the structure thin pili of *A. baylyi* BD413 consists of multiple hydrophobic interactions between several filaments which makes it a hydrophobic appendage that promotes binding to hydrophobic surfaces [36]. Presence of thin pili in *A. baylyi* BD413 suggests that they play a significant role in the interaction/repulsion between bacteria and hydrophobic/hydrophilic surfaces.

Biosurfactants including lipopeptides are recognized as a main factor in microbial attachment to solid surfaces [37]. It has been demonstrated that *Bacillus* spp. can produce cyclic lipopeptides such as surfactin, iturin, fengycin and lichenisin to confer cell surface hydrophobicity (CSH) to the strains [38]. CSH is involved in adhesion to hydrophobic surfaces and is a plausible factor in *Bacillus* attraction towards hydrophobic substrates. Kesel et al. also suggested that the thiol groups present in bacterial cell membrane or biofilm matrix proteins expressed at low levels at the beginning of *B. subtilis* NCIB 3610 adhesion to hydrophobic gold surface, mediates the initial attachment [39].

Negatively charged surface of both bacterial cells [40, 41] also explains bacterial adsorption towards positively charged surfaces with NH_2_^+^ and repel from negatively charged surfaces with COO^-^.

SEM images of *B. subtilis* biofilms on CH_3_ and NH_2_^+^ shows that among vegetative cells inside the biofilm structure, spores are present. Branda et al. proved that sporulation happens during the biofilm formation process [42] and Lindsay et al. suggested that when exposed to nutrient limitation stress, cells inside the biofilms of *B. subtilis* may be stimulated to form spores [43]. Sporulation and matrix production are linked together via Spo0A signaling pathway [44] and can explain why we were able to detect both forms of bacterial life on the surfaces. SEM images of biofilms on CH_3_ show higher numbers of spores compared to NH_2_^+^, which might be because the fact that spores of each bacterial cell are more hydrophobic than vegetative cells [41] and so more likely to interact with hydrophobic surfaces instead of hydrophilic substrate with NH_2_^+^.

Hydrophilic surface with a positive surface potential affects *A. baylyi* cell morphology to be elongated which can be because of a better distribution of intrinsically polar EPS. Interestingly, the lysis of the bacterial cell wall was observed with a positive zeta potential for both bacterial strains due to the strong electrostatic interactions (see Fig. S6 in supplementary documents).

Our study was conducted for 72h started in a nutritive condition that allowed us to monitor biofilm formation stages on different surface types. Metabolic activity results showed that the number of adhered cells increased from the beginning of incubation until 24h when biofilm reached the maturation apex. After the first step of rise, decrease in metabolic activity was observed likely due to the detachment of cells or parts of the biofilm. Metabolic activity reduction was continued until 72h with no further increase for all the solid substrates, proving the same cyclical behavior of biofilm for both bacterial strains on all the substrates. Morgenroth and Wilderer, 2000 reported successive increases and decreases in biofilm cycles formed under flow conditions with no respect to the effect of surface characteristics [45]. Ploux et al. demonstrated that cyclical behavior of *E. coli* biofilm is dependent on material surface characteristics on substrates with NH_2_^+^ and CH_3_ [31]. Their experiments were done using a minimal medium and a long duration of incubation time (336h). Our study also adds the results of biofilm formation on hydrophilic negatively charged COO^-^ and neutral charged OH surfaces in highly nutritive media for a shorter time duration (72h) for two different bacterial strains. Results show that the cycle duration not only depends on the wettability, but also the surface charges. The shortest cycle duration was recorded for negatively charged hydrophilic surfaces with COO^-^ for both bacterial strains while the longest was captured for hydrophobic surfaces. The OH and COO^-^-substrates showed neither support for bacterial initial attachment nor preserving them from detachment. As it can be noticed on SEM images, only scattered bacterial cells attached to the OH and COO^-^ could be observed. These we assume can easily detach from the surface compared to a well-established biofilm. The difference between the structure of biofilms on NH_2_^+^ and CH_3_ also confirms that in *A. baylyi* monolayer biofilm and in *B. subtilis* tree-like (filamentous) biofilm on NH_2_^+^, bacteria were more isolated than in multilayer biofilms on CH_3_. Bacteria are less protected by their neighbors and are more prone to be detached on surfaces with NH_2_^+^ than on CH_3_-substrates. Logically, different substrate characteristics lead to different environmental conditions followed by different kinetics of biofilm attachment and detachment cycle durations.

Studies investigating biofilm formation on controlled surfaces mostly have focused only on the microbial cell distribution and arrangement structure but ignored the EPS production changes in response to the surface characteristic. In our study, changes in biochemical composition of the biofilm matrix in response to different surfaces in nutritive media were monitored over time intervals using micro-FTIR. The matrix of *B. subtilis* is mainly composed of TasA protein [46], biofilm surface layer protein A (BslA) protein [47, 48]), polysaccharides [42] and eDNA [49]. TasA, a functional amyloid, is the main component of the biofilms of *B. subtilis* and supports the scaffold of biofilms by its fiber structure. Although Diehl et al. demonstrated that TasA has heterogeneity in its secondary derivatives, robust fibers enriched in β-sheets have been detected as the predominant conformation of TasA in biofilms. Micro-FTIR results prove β-sheets secondary structures for *B. subtilis* biofilms formed on hydrophobic surfaces with CH_3_ and hydrophilic surfaces with NH_2_^+^, which we assume to be from TasA. BslA is a self-assembling secreted protein by *B. subtilis* that coats biofilm surface and renders hydrophobic features and water repellency. TasA is essential for initial attachment and colonization of *B. subtilis* in the rhizosphere, demonstrating its importance in bacteria-surface interactions and clearly shows the need for in-depth investigation into its role in bacterial attachment to different surfaces. BslA also has a contribution in biofilm matrix assembly by interacting with TasA and exopolysaccharides. X-Ray diffraction analysis of BslA shows the architecture of β-sandwiches consist of 310 helix and β- strands. To the best of our knowledge, there is no comprehensive evidence about the matrix composition of *A. baylyi* biofilms. Gregorio, et al., reported that Biofilm associated protein (Bap) proteins are conserved among *Acinetobacter* spp. [50], so it is possible to speculate that Bap plays a major role in *A. baylyi* biofilm matrix as they do in *Acinetobacter baumannii* biofilms. Other proteins that are assumed to be involved in *A. baylyi* biofilm formation are fimbrial-biogenesis protein (3317) and putative surface protein (PEVL 389) [51]. Our FTIR results show that at the matured biofilms, proteins are the dominant composition of the matrix on the substrates tested. Also peaks for DNA, phospholipids and polysaccharides were detected on 24h biofilms. After 6h of exposure, protein signature was observed in the IR spectra for both strains which are possibly from the attaching proteins (pili for *A. baylyi* and lipopeptides of *B. subtilis*). Spectra shows that protein production was constant over the 3 times. Variations in the appearance of the bands assigned for polysaccharides, DNA, and lipid show that they were not necessary for the initial biofilm formation but for maturation and detachment. Secondary derivative and amide I/amide II analysis testify the higher colonization potential of both strains to CH_3_ and NH2+. The lowest absorbance intensity and the lowest content of amide I/amide II was recorded for both bacterial strains on COO-, while *B. subtilis* showed higher affinity towards OH than *A. baylyi*.

The use of functionalized surfaces as a physiochemical tool to manipulate surfaces toward preservation from biofilms or promotion of biofilms is a promising approach. With the analysis of chemical, physiology, morphology and compositional changes, the present study shows how bacterial biofilms are affected by different surface characteristics as a function of time. To control negative biofilms, hydrophilic surfaces with COO^-^ and OH seem to be the surface of choice as they do not support bacterial attachment and biofilm formation. However, the highest number of bacterial cell attachment and matrix production on hydrophobic solid surfaces with CH_3_ is a very interesting concept for rewilding and forming positive biofilms wherever are needed. Positive biofilm formation on desired surfaces is an intriguing field to explore in future work.

## Conclusion

Our work indicates that there is a distinct pattern of cells being most attracted to hydrophobic or positively charged substrates, while a lack of attachment and biofilm formation on hydrophilic and negatively charged surfaces. There are minor variations between the two bacterial species tested, but in general, it seems to be a universal pattern across the Gram-barrier. Comparison of the two attractive surfaces reveal that substrate variation does not seem to induce a noticeable change in temporal matrix composition, but rather favor accumulation of biomass and successful biofilm development. On a general note, these results highlight the potential of surface treatments in areas where it is intended to either repel or attract bacteria.

## Supporting information

supplementary doc

## Acknowledgements

The authors would like to thank Jeremiah Shuster, Klaus Qvortrup and Cristiano di Benedetto for their help in SEM sample preparation and imaging. The contributions from SH, IK, TV, and KKS was supported by a research grant from VILLUM FONDEN (00025352)

## Material and methods

### Substrate preparation

In this work, four functionalized self-assembled monolayers with OH, COO^-^, NH_2_^+^ and CH_3_ were prepared. Gold Coated Ø4 ‘‘ Silicon wafers, 50 nm Au on 500μm P-type were purchased from tedpella and were cut to chips with the dimensions of 10 mm × 10 mm. The chips were cleaned by washing three times with ethanol, then left in an UV ozone cleaner for 20 min, followed by rinsing with ethanol. Thiol solutions were prepared as table 2 and stored at 4 up to 6 months. To prepare the SAMs on gold surfaces, the cleaned chips were immediately soaked in thiol solutions for 18h at room temperature. After incubation, chips were rinsed with ethanol and sterile water.

**Table 2.** SAMS thiol solution preparation.

| SAMs | Chemical | Powder (mg) | Solvent (ml) |
| --- | --- | --- | --- |
| CH<sub>3</sub> | 1-undecanethiol | 19 | 100 ml ethanol* |
| NH<sub>2</sub><sup>+</sup> | 1-amino-1-undecanthiol hydrochloride | 24 | 100 ml ethanol |
| COO<sup>-</sup> | 1-Mercaptoundecanoic acid | 22 | 90 ml ethanol + 10 ml acetic acid |
| OH | 1-Mercapto-1-undecanol | 20 | 100 ml ethanol |

\*Ethanol used in the preparation was absolute 200% ethanol

### Bacterial strains and culture conditions

The bacterial species *B. subtilis* 168 and *A. baylyi* BD413 were used in this work. The strains were first cultivated in separate pure cultures overnight on Tryptic soy agar (TSA; Merck, Germany) at 37°C. Several colonies were used to inoculate 25 ml of Tryptic soy broth (TSB; Merck, Germany) for *B. subtilis* and Loria berthany (LB; Merck, Germany) for *A. baylyi*. These initial cultures were incubated at 37°C with shaking at 180 rpm for 18 h corresponding to ~10^8^ cells. The cells were centrifuged at 5000 rpm at 4°C for 10 mins and resuspended in prechilled LB containing 15% glycerol, aliquoted and stored at −80°C until further use.

### Attachment and biofilm formation experiment

The biofilm-forming ability of *B. subtilis* and *A. baylyi* was evaluated in a static model system. SAMs substrate chips were aseptically placed in 48-well sterile flat-bottomed microtiter plates using sterile tweezers. 1.5 ml of overnight bacterial culture with OD600 adjusted to 0.1 was added to each well. The strain was grown on the substrates at 30 °C. Several culture durations were explored: 6h, 12h, 24h, 36h, 48h, and 72h. One series of six CH_3_, six OH, six COO^-^, and six NH_2_^+^ -substrates were prepared for each culture duration. After incubation, to eliminate non-adhered or loosely adhered bacterial cells, test samples were rinsed twice, by immersion in fresh sterile 0.9% saline. The metabolic activity of attached cells was determined using triphenyl tetrazolium chloride (TTC). TTC solutions in distilled water were prepared by dissolving TTC powder (Sigma-Aldrich) in distilled water in concentration of 0.5% and then sterilized by 0.22 μm cellulose acetate filters. Chips were transferred to a new sterile 24-well plate. 300 μL of TTC solution and 1200 μl of LB or TSB were added to all the wells for *A. baylyi* and *B. subtilis*, respectively. Plates were incubated in the dark for 24 h at 37 °C. After the incubation period, 200 μl of the well contents were moved to a new 96-well flat-bottomed microplate and the absorbance was measured at 405 nm using XXX.

### Biofilm structure morphology

Morphology of biofilms formed on SAMS substrates was visualized using both scanning electron microscopy (SEM) and confocal laser scanning microscopy (CLSM). For each type of bacteria-substrate combination two samples were prepared after 24 h of incubation for SEM and 3h, 24h and 48h for CLSM. For each sample, several places were focused and a series of images was taken by systematically displacing the samples under the camera.

### SEM imaging

Biofilms were formed on 10 mm × 10 mm chips for 24h and after rinsing twice in fresh NaCl solution (9 g/l in water), test samples were prepared following the classical steps: samples were chemically fixed by immersing overnight in a 4% glutaraldehyde (w/v in 0.1 PBS) solution, dehydrated by immersing in several ethanol solutions (from 20% (v/v) in water to 100%). Samples were dried using a critical point dryer (Leica (Balzer) CPD030). Finally, samples got sputter-coated with gold (Leica Coater ACE 200) at room temperature.

Examination was done at several magnifications (from 5000× to 100,000×) using a versatile high-resolution dual beam SEM (FEI Quanta 3D FEG) with high vacuum mode, Everhart-Thornley Detector(ETD).

### CLSM imaging

For microscopic investigation of bacterial attachment and biofilm formation of surfaces with different properties, cells were exposed to Custom-made gold chips for 3, 24 and 48h. patches were stained with FilmTracer LIVE/DEAD biofilm viability kit (Thermo Fisher) with SYTO9 at a final concentration at 10μM and Propidium Iodide at 60 μM. The concentration of Fluorescent stains were adjusted with sterile 0.9% NaCl. Patches were stained for 20 min in darkness, before Images were acquired on a single point confocal laser scanning microscope (Zeiss LSM800, Carl Zeiss Inc. Germany) equipped with an EC Plan-Neofluar x20/0.5 air objective and an Axiocam 503 mono camera. For single samples where higher magnification was required a Plan-Apochromat x63/1.4 oil objective was used. Syto9 stained cells were emitted using a 488nm laser, while the propidium iodide stain was emitted with a 561 nm laser. All Images were 1024×1024 pixels (0.312 um/px for 20x and 0.105um/px for 63x) and when 3d images were required an interval of 0.8um was used between Z-stacks.

### Micro-FTIR

The micro-FTIR spectra of the biofilms grown on different substrates were recorded in transflection mode using Hyperion 3000 IR microscope coupled with a Tensor 27 FTIR spectrometer (Bruker Optik, Ettlingen, Germany) and equipped with a globar light source and KBr beamsplitter. The microscope is further equipped with a single-element liquid-nitrogen-cooled mercury cadmium telluride (MCT) detector and a ×15/0.4 objective. Spectra in the range of 600 - 3900 cm^-1^ were recorded using a spectral resolution of 4 cm^-1^. For one resultant spectrum (*I*), 256 interferograms were averaged and the result was Fourier transformed into a spectrum applying the BlackmannHarris 3 apodization function and zero-filling factor 2. Before the measurements, the biofilms grown on SAMS chips for 24h were washed twice by flooding the chips with sterile deionized pure water and air dried at room temperature overnight. The background spectra (*I_0_*) were recorded using the same parameters on biofilm-free areas of the substrate with medium and reflectance spectra (*R*) were then obtained as:

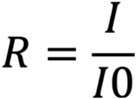

Recorded reflectance spectra were converted to absorbance followed by a correction to remove spectral bands of atmospheric CO_2_ and water vapor (OPUS software, Bruker). Spectral range between 600 and 1800 cm^-1^ (fingerprint region) was selected for further analysis. Baseline correction was then applied with baseline points at 1775, 1480, 1358, 1185, 804 and 603cm^-1^.

### Statistical analysis

All experiments were performed in triplicates and values were expressed as means ± standard deviation (SD). Comparisons between means of groups were analyzed using one-way ANOVA and paired t-test using SPSS 18. P < 0.05 was considered statistically significant and * represents p<0.05; ** p<0.01; *** p<0.001 and **** p<0.0001.

