## supplementary doc for "Fundamentals of biofilm formation in soil: From functionalized self-assembled monolayers to rewilding"

Appendix: **Supplementary information**

**X-ray Photoelectron Spectroscopy (XPS) analyses of SAMs preconditioned in nutrient media.**

Supplemental Methodology

SAMs chips were prepared as explained in materials and methods and were aseptically placed in 48-well sterile microtiter plates using sterile tweezers. 1.5 ml of one of the three media, tryptic soy broth (TSB), Luria Broth (LB) and minimal media 9 (M9), were added to each well. Plates were incubated overnight under static conditions at 37. Samples were rinsed twice with sterile molecular grade water and air dried overnight at room temperature. As a control, SAMs not incubated in media were used.

The surface chemistry of the various wafer pieces (one replica) was analyzed using a Kratos Axis Ultra DLD spectrometer using a monochromatic Al K alpha source (1486.6 eV) operated at 150W. For survey spectra a pass energy of 160 eV and a step size of 1 eV was used and for high resolution spectra 20 eV with a step size of 0.1 eV. A hybrid lens system with a magnetic lens provided an analysis area of 0.3 x 0.7 mm. The built-in charge neutralizing equipment was used to compensate for sample charging and the binding energy scale was calibrated by referencing the aliphatic carbon peak at 285.0 eV. All measurements were done at room temperature. In general, the error in XPS analyses is estimated to be less than 10%. Quantification was done using CasaXPS with peak shapes GL(30) on high resolution spectra.

Supplemental Results and Discussion

The four control surfaces all show presence of the expected functional groups in the spectra. The quantity of thiols from the anchoring functionality was found to be around 2 at % for all SAMS except the $NH_2$ - terminated SAM that showed 1.3 at% (Table 1). The oxygen present in the samples with SAM-$NH_2$ may come from the underlying substrate $SiO_2$, under the Au layer, or is a result of impurities.

After exposure to culture medium an overlayer is formed which can be seen in the decrease of the signal from the substrate for all exposed samples. This is seen as a decrease of the underlying Au signal and also a decrease in the thiol positioned at the bottom of the SAM. In parallel to this, there is a small increase in the total quantity of C and N for the incubated samples. For SAM- OH or SAM-CH3 also a small increase in total content of O was observed. For SAM- OH or SAM-COOH a small quantity of Na was also present in all treated samples (except SAM-OH incubated in LB). For SAM-OH this is significant as the SAM does not contain N, the same if true for SAM-COOH that does not contain Na, and SAM-CH3 that does not contain either O or N when prepared. The changes in the SAM-NH2 is within the estimated 10% error of XPS and therefore uncertain. However, all the observed increases are small. Therefore, they do not allow for ranking of efficiency of preconditioning between the four SAMS. In order to do so a much larger number of replicas would have needed to be analyzed. However, the data indicate that low levels of substances are adsorbed from the various media onto the SAMs. The quantity is too low to make any hypothesis of what they may be, except that Na most likely is balancing negatively charged groups on the SAM surface. Survey spectra of the samples can be found as Figures S1-S4.

**Table S1**. XPS data showing low levels of preconditioning by growth medium on functionalized surfaces made from self-assembled monolayers (SAM) terminated with OH, $COO^-$, $NH_2^+$ and $CH_3$. The investigated growth media were tryptic soy Broth (TSB), minimal media 9 (M9), and Luria Broth (LB).

| **elemental atomic % of surface** | **Na 1s** | **O 1s** | **N 1s** | **C 1s** | **S 2p** | **Au 4f** |
|---|---|---|---|---|---|---|
| SAM-OH | | 5.9 | | 47 | 2.4 | 45 |
| SAM-OH + TSB | 0.9 | 8.1 | 1.6 | 51 | 1.6 | 36 |
| SAM-OH + M9 | 2.2 | 7.3 | 1.8 | 55 | 1.6 | 32 |
| SAM-OH + LB | | 7.1 | 0.7 | 54 | 1.8 | 36 |
| SAM-COOH | | 11 | | 50 | 2.0 | 37 |
| SAM-COOH + TSB | 0.7 | 11 | 4.4 | 58 | 1.5 | 24 |
| SAM-COOH + M9 | 1.9 | 11 | 2.1 | 53 | 1.7 | 30 |
| SAM-COOH + LB | 1.5 | 11 | 6.6 | 59 | 1.5 | 20 |
| SAM-NH2 | | 6.4 | 3.6 | 65 | 1.3 | 23 |
| SAM-NH2 + TSB | | 7.0 | 4.7 | 71 | 0.9 | 17 |
| SAM-NH2 + M9 | | 5.5 | 2.7 | 73 | 0.9 | 18 |
| SAM-NH2 + LB | | 9.1 | 4.8 | 73 | 0.7 | 12 |
| SAM-CH3 | | | | 45 | 2.1 | 53 |
| SAM-CH3 + TSB | | 3.8 | 2.3 | 50 | 1.9 | 42 |
| SAM-CH3 + M9 | | 6.3 | 2.0 | 46 | 1.6 | 44 |
| SAM-CH3 + LB | | 3.9 | 2.8 | 48 | 2.1 | 43 |

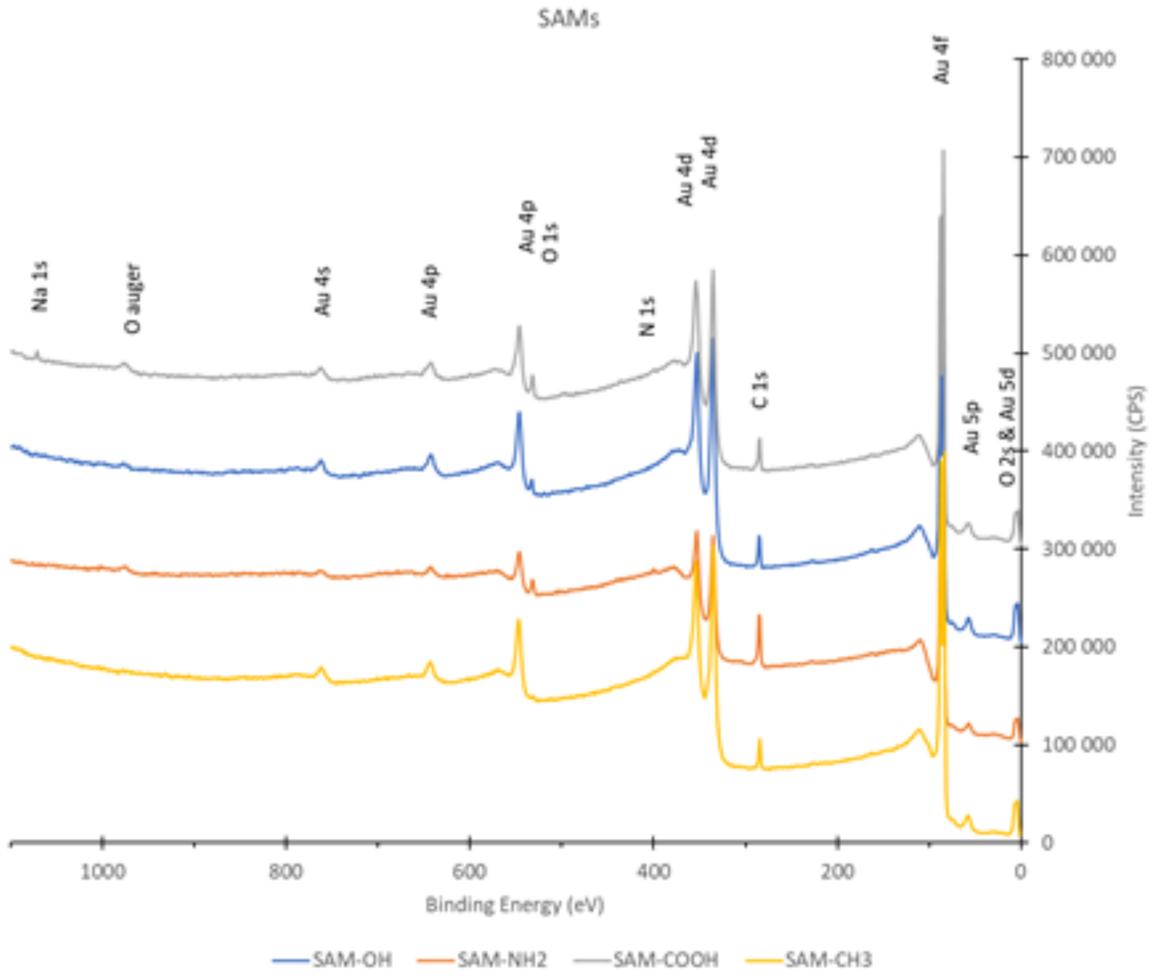


Figure S1. XPS survey spectra of self-assembled monolayers prepared on Au coated Si wafers. Grey line represents $COO^{-}$-terminated SAM, blue OH-terminated SAM, orange $NH_2^{+}$-terminated SAM and yellow $CH_3$-terminated SAM.

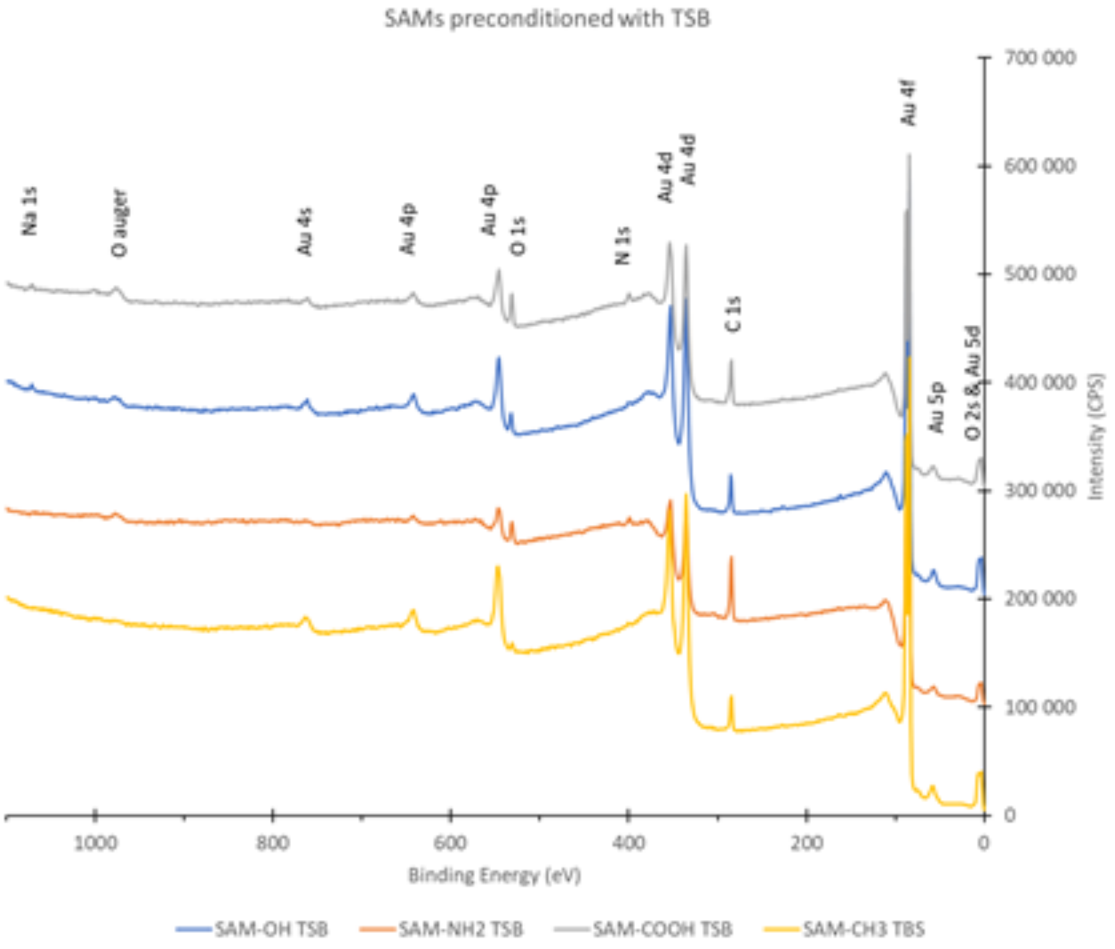


Figure S2. XPS survey spectra of self-assembled monolayers prepared on Au coated Si wafers and preconditioned in TSB medium. Grey line represents $COO^-$-terminated SAM, blue OH-terminated SAM, orange $NH_2^+$-terminated SAM and yellow $CH_3$-terminated SAM.

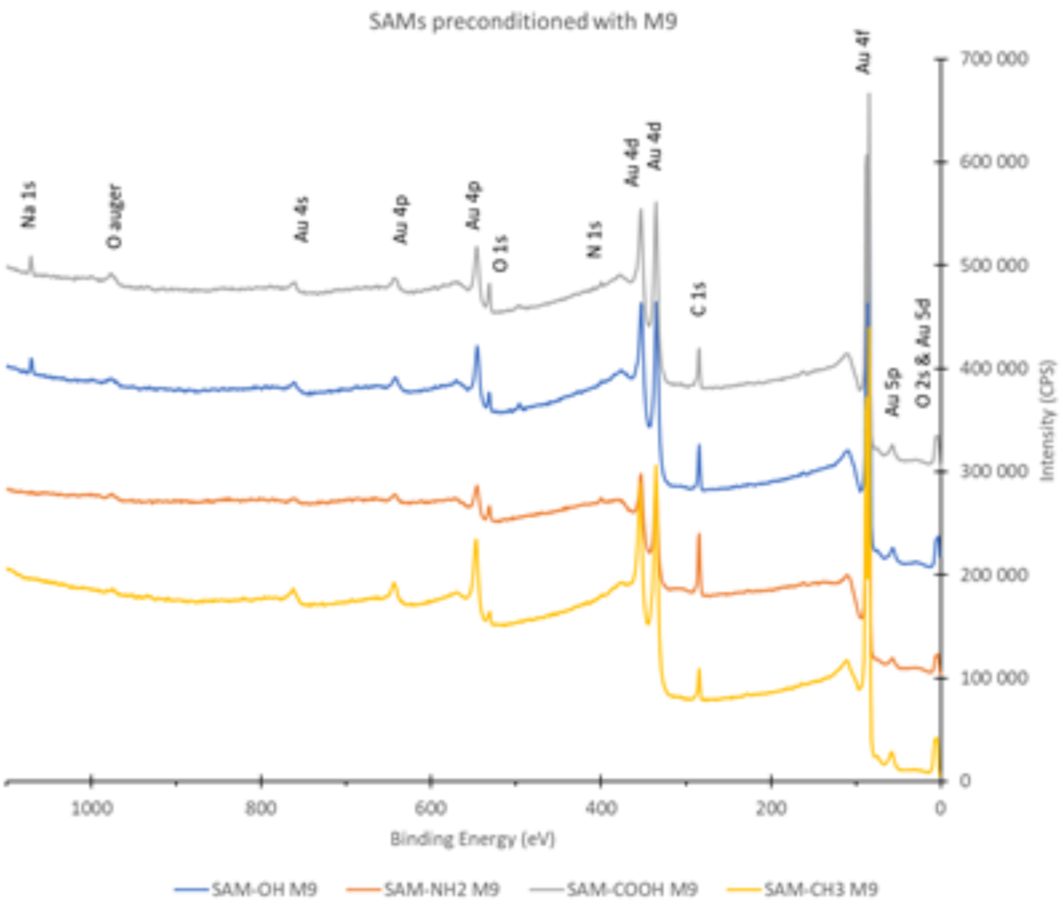


Figure S3. XPS survey spectra of self-assembled monolayers prepared on Au coated Si wafers and preconditioned in M9 medium. Grey line represents $COO^-$-terminated SAM, blue OH-terminated SAM, orange $NH_2^+$-terminated SAM and yellow $CH_3$-terminated SAM.

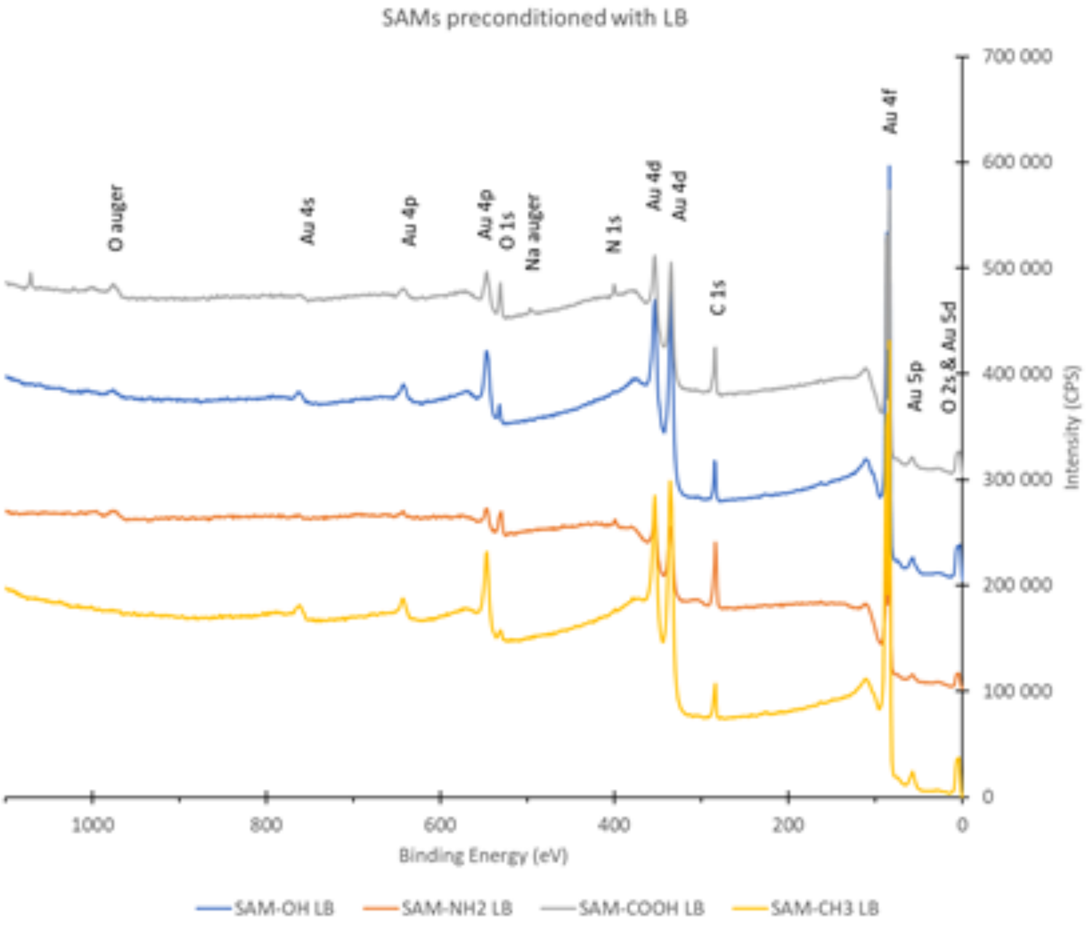


Figure S4. XPS survey spectra of self-assembled monolayers prepared on Au coated Si wafers and preconditioned in LB medium. Grey line represents $COO^-$-terminated SAM, blue OH-terminated SAM, orange $NH_2^+$-terminated SAM and yellow $CH_3$-terminated SAM.

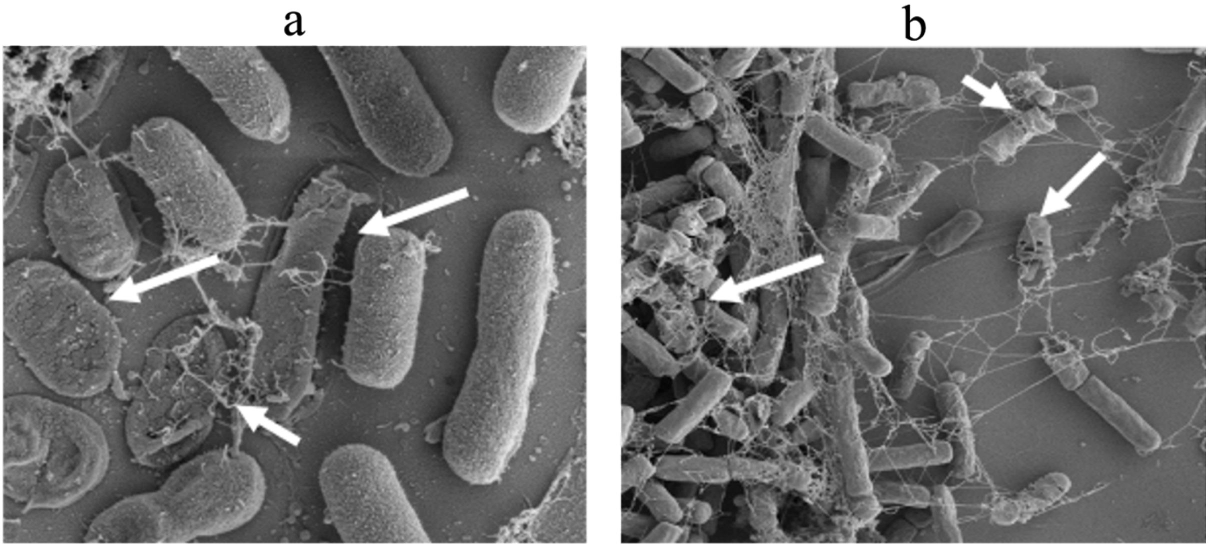


Fig S5. Bacterial cell lysis observed by SEM. a) *A. baylyi*, b) *B. subtilis*. Arrows show lysed cells.

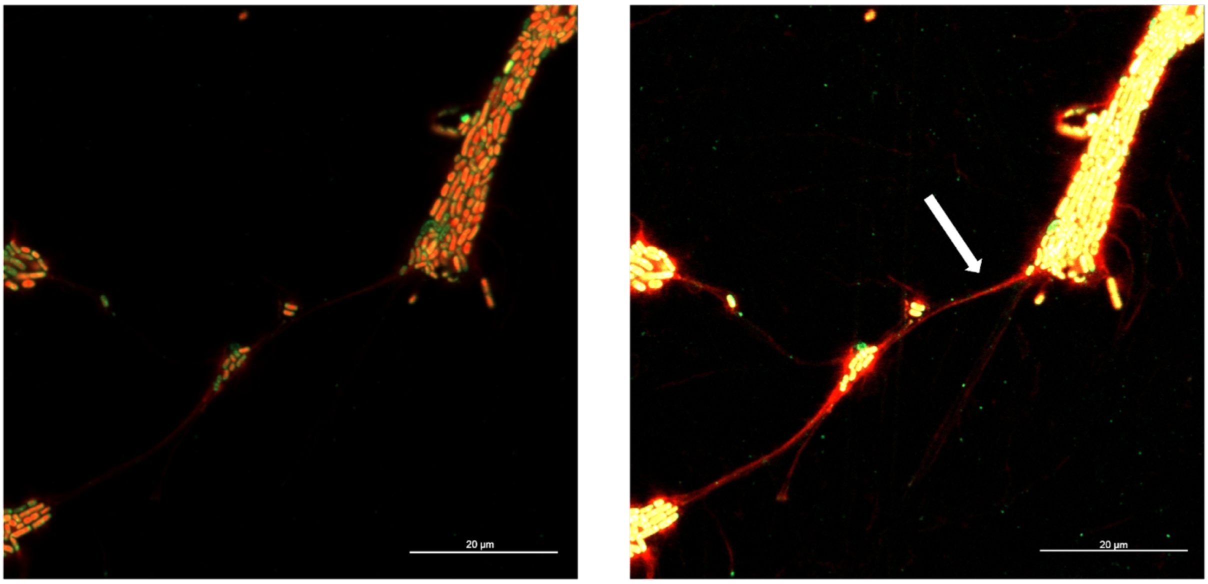


Fig S6. Images obtained by CLSM (63X) after LIVE/DEAD biofilm viability kit staining with two contrasts. SYTO9 stained alive cells green and PI stained dead cells and eDNA red. Arrow shows eDNA string around biofilm.
